# Long-term antiplatelet therapy protects against cerebral but not parenchymal amyloid plaque formation and neurodegeneration in transgenic mice of Alzheimer’s disease

**DOI:** 10.1101/2025.06.30.662310

**Authors:** Lili Donner, Laura Mara Toska, Marlene Pils, Fabian Rehn, Oliver Bannach, Margitta Elvers

## Abstract

**Introduction:** Cerebral amyloid angiopathy (CAA) is characterized by the aggregation of amyloid-β peptides in cerebral blood vessels, leading to a loss of vascular integrity and contributing to the progression of Alzheimer’s disease (AD). In our previous study, we showed that short-term treatment with the antiplatelet drug clopidogrel, a P2Y12 inhibitor that irreversibly blocks ADP signaling, reduced the incidence of CAA in the APP23 mouse model of Alzheimer’s disease providing strong evidence for platelets to contribute to AD pathology. The objective of the present study was to ascertain whether long-term treatment with clopidogrel and the earlier initiation of treatment prior to the formation of Aβ deposits prevents pathological changes associated with Alzheimer’s disease.

**Methods:** APP23 mice were treated with clopidogrel for 15 month and analyzed for AD pathology.

**Results:** We detected increased permeability of the blood brain barrier in different brains regions of APP23 mice. Moreover, platelets migrated into the brain parenchyma and accumulated around amyloid plaques in the brain of APP23 mice. Although we detected platelets in close proximity to microglia and neurons in the cortex and hippocampus of APP23 mice, we did not observe any differences in neurodegeneration or gliosis in Clopidogrel treated APP23 mice. Furthermore, pathological analysis showed a significant reduction in CAA and in soluble Aβ42 levels in the brain of Clopidogrel versus placebo treated APP23 mice but no differences in plaque load in the brain parenchyma.

**Conclusion:** Thus, antiplatelet therapy may alleviate amyloid pathology in cerebral vessels leading to improved blood perfusion in AD patients.

## 1 Introduction

Around 57 million people worldwide suffer from dementia. Since the age of the general population increases alongside with lack of effective therapeutic strategies, the number of people suffering from dementia has rapidly increased, estimated to reach 152 million by 2050 (2022). The most common form of dementia, which causes a progressive decline in cognitive function, is Alzheimer’s disease (AD) (Clarfield, 2003). The histopathological hallmarks of AD include the accumulation and aggregation of amyloid-β (Aβ) in the brain parenchyma as amyloid plaques and in the cerebral vasculature known as cerebral amyloid angiopathy (CAA), the formation of intracellular tau tangles, and neuroinflammation leading to neuronal loss (Selkoe, 2011, Thal et al., 2008). Approximately 80% of Alzheimer’s disease patients exhibit CAA, which leads to significant dysfunction of the cerebral vasculature and breakdown of the blood-brain barrier (BBB) (Thal et al., 2008).

Platelets, the smallest cells in the blood, are not only involved in the regulating of hemostasis and thrombosis, but also modulate processes of acute and chronic inflammation in disease pathology (Koupenova et al., 2018). In addition, platelet dysfunction is associated with several neurodegenerative diseases, including Alzheimer’s and Parkinson’s disease (Leiter and Walker, 2020, Beura et al., 2022). Platelets are thought to be the main source of Aβ peptides and amyloid precursor protein (APP) in the circulation and can release them during activation (Leiter and Walker, 2020, Wolska et al., 2023, Gowert et al., 2014, Li et al., 1998, Chen et al., 1995). Analyses of aged transgenic mice modeling Alzheimer’s disease (APP23) with parenchymal plaques and CAA have shown that platelets isolated from these mice are in a pre-activated state in the circulation and adhere to vascular amyloid-β deposits, leading to cerebral vessel occlusion (Jarre et al., 2014, Gowert et al., 2014).

In our previous studies, we found that platelets are able to convert soluble Aβ40 peptides into fibrillary Aβ aggregates *in vitro* (Gowert et al., 2014, Donner et al., 2020, Donner et al., 2018, Donner et al., 2021, Donner et al., 2016). Moreover, we identified two direct binding partners of Aβ40 at the platelet surface, the fibrinogen receptor integrin αIIbβ3 and the major collagen receptor glycoprotein VI (GPVI). The blocking or deletion of these receptors led to a significant reduction of Aβ aggregation *in vitro* (Donner et al., 2016, Donner et al., 2020). We also found that ADP plays a role in platelet-mediated Aβ aggregation, as inhibition of the ADP receptor P2Y_12_ also prevented Aβ aggregation *in vitro* (Donner et al., 2016). In addition, we have shown that a short-term treatment with the antithrombotic drug clopidogrel, a P2Y_12_ antagonist, for three months resulted in a reduction of CAA in APP23 mice (Donner et al., 2016). However, it is unclear whether treatment before the onset of pathology and long-term treatment with clopidogrel leads to the prevention of AD pathology in transgenic APP23 mice.

Therefore, we treated APP23 mice with the antiplatelet agent clopidogrel for 15 month starting at 2 months of age, before the mice develop amyloid-β pathology. We detected a significant reduction in the level of soluble Aβ42 in brain lysates and reduced CAA in clopidogrel treated compared to untreated APP23 mice. However, the plaque load in the brain parenchyma, gliosis and neuronal loss were unaffected by clopidogrel treatment.

## 2 Materials and Methods

### 2.1 Animals

The transgenic APP23 AD mouse model on C57BL/6J background, introduced by Sturchler-Pierrat (Sturchler-Pierrat et al., 1997) in 1997, was used. At 3 months of age develop these mice cognitive deficits, which are detectable in the Morris water maze and at 6 months develop amyloid plaques (Van Dam et al., 2003). The Aβ pathology and cognitive deficits progressively intensify with age (Van Dam et al., 2003, Sturchler-Pierrat and Staufenbiel, 2000). Food and water were provided ad libitum. Control groups (male heterozygous APP23 mice (placebo) and their male non-transgenic littermates (WT)) were fed standard chow, and male APP23 mice were fed standard chow supplemented with clopidogrel (Actavis). Based on the amount of food consumed per day, the daily intake of clopidogrel was approximately 2 mg. The average age at the start of treatment was 2 months ± 2 weeks. After 15 months of treatment the mice were sacrificed. The right hemisphere was snap frozen in liquid nitrogen for determination of Aβ peptides and the left hemisphere was fixed in 4 % paraformaldehyde (PFA) overnight and then in 30% sucrose for two days and snap frozen in cold isopentane for histopathological analysis.

To analyse platelet migration into the brain parenchyma, the global double-fluorescent Cre reporter mouse mT/mG, which expresses membrane-targeted tdTomato before Cre excision and membrane-targeted EGFP after Cre excision, was crossed with the megakaryocyte/platelet-specific PF4-Cre transgenic mouse to confine the expression of EGFP to megakaryocytes and platelets. These mice were crossed with heterozygous APP23 mice to generate the APP23mT/mG;PF4Cre+; and the WTmT/mG;PF4Cre+ mice. 6, 16 and 24 month-old APP23mT/mG;PF4Cre+ and WTmT/mG;PF4Cre+ mice were anaesthetised intraperitoneally with ketamine (100 mg/kg) and xylazin (10 mg/kg) and then transcardially perfused first with ice-cold 0.9% NaCl and then with ice-cold 4% PFA. The brains were removed and stored in 4% PFA overnight. The next day, the brains were stored in 30% sucrose for two days and finally snap frozen in cold isopentane.

All experiments were performed in accordance with the German Law on the protection of animals and approved by a local ethics committee (LANUV, North-Rhine-Westphalia, Germany, reference number: reference numbers O 86/12, AZ 84-02.05.40.16.073, AZ 81-02.05.40.21.041, AZ 81-02.04.2019.A232).

### 2.2 Immunofluorescence staining

IF was performed on 14-µm thick coronal sections containing hippocampus frozen brain sections. Five to six slices of each mouse were stained for each IF. In brief, the sections were washed three times for 10 min with PBS. Sections were blocked with 5 % goat serum and permeabilized with 0.3% Triton X100 (Sigma) for 1-2 hours at RT, then washed three times with PBS. Sections were incubated with the primary antibody (anti-β-Amyloid (6E10), 1:200, BioLegend; anti-Aβ40 (11A50-B10), 10 µg/ml, BioLegend; anti-Aβ42 (12F4), 1 µg/ml, BioLegend; NeuN, 1:1000, Abcam; Iba1 (JM36-62), 1:100, Invitrogen; GFAP (GA5), 1:100, Invitrogen; CD31 (550274), 1:75, BD Biosciences), or their isotype control immunoglobulins in blocking solution (1 % goat serum and 0.3 % Triton X100) over night at 4 °C in a humid chamber. After three washes with PBS, brain sections were incubated with fluorochrome-conjugated secondary Abs from Invitrogen (Goat anti-Mouse IgG Alexa Fluor™ 660; Goat anti-Rat IgG Alexa Fluor 647; Goat anti-Rabbit IgG Alexa Fluor 647; Goat anti-mouse IgG Alexa Fluor® 488; Goat anti-Rabbit IgG Alexa Fluor™ 555) at RT for 1 h. For IgG staining, brain slices were blocked with blocking solution (3 % BSA with 0.1 % Triton in PBS) at RT for 2 h, incubated with Goat Anti-Mouse IgG Antibody (H+L), Biotinylated (Vector,1:150) afterwards used as secondary antibody Streptavidin-660 (Vector, 1:50). Nuclei were stained with 4′,6-Diamidine-2′-phenylindole dihydrochloride (DAPI, Roche, 1:3000) for 5 min. Immunofluorescence images were acquired with the AX10 Observer.D1 HAL100 inverted phase-contrast fluorescence microscope (Carl Zeiss, Oberkochem, Germany). To quantify the genetically expressed GFP signal of the APP23mT/mG;PF4Cre+; and the WTmT/mG;PF4Cre+ mice, the mean fluorescence intensity (MFI) value was measured in the cortex and hippocampus regions of the brain with the same exposure time. The quantification was conducted at 100 x magnification. The quantification of immunoglobulin G, GFAP and Iba1 was conducted using the MFI value with an identical exposure time for all sections. The evaluation of MFI was carried out via the program Zen2.6 lite blue. The area fraction and plaque number of 6E10-positive staining in cortex and hippocampus and Aβ40- and Aβ42-positive vessels were quantified with ImageJ software (National Institute of Health, Bethesda, USA). Images of all the areas with Aβ40 or Aβ42 positive vessels were acquired, thresholded using ImageJ and the total Aβ-positive area per section was calculated. To ensure comparability between sections, the threshold was maintained across images of the same staining. Three to five sections of each mouse were stained for each immunfluorescence staining. The cortex region was analysed with three images per section, while the hippocampus region was analysed with one or two images (whole hippocampus). The counting of double-positive DAPI and NeuN cells in the CA1 region was performed using QuPath software (University of Edinburgh, UK).

### 2.3 Congo red staining

Cerebral amyloid angiography (CAA) was visualized by Congo red staining. Sections were washed with water for 10 min and then stained with 1% Congo red dye (Merck) solution at RT for 10 min. The sections were then immersed in 1% lithium carbonate for 15 seconds and then in 80% ethanol. Finally, the sections were stained with Mayer’s hematoxylin (Carl Roth) for 3 min and washed with tap water for 10 min. After washing, the sections were immersed in 70%, 96% and 100% ethanol for one minute each. After the ethanol series, the sections were immersed in Roti®-Histol (Carl Roth) for 5 min and then mounted with Roti®-Histokitt (Carl Roth). Four to five slices of each mouse were stained and visualized under fluorescent microscope (AX10 Observer.D1 HAL100 inverted phase-contrast fluorescence microscope, Carl Zeiss, Oberkochem, Germany). CAA number per slice were quantified.

### 2.4 Prussian blue staining

Prussian blue staining was performed to detect the presence of cerebral haemorrhages. Sections were incubated with 5% potassium hexacyanoferrate(II) trihydrate for 5 min and then with freshly prepared 5% potassium hexacyanoferrate trihydrate and 5% hydrochloric acid for 30 min. After rinsing in water, sections were counterstained with Nuclear Fast Red for 10 min and then rinsed in water. Finally, the sections were dehydrated and Roti®-Histokitt was used as a mounting medium. Hemosiderin deposits were identified at a 20× and 40× magnification.

### 2.5 Measurement of soluble and insoluble Aβ40 and 42 peptides

The amount of soluble and insoluble Aβ40 and Aβ42 in the brains was determined by performing a four-step ultracentrifugation. The right hemispheres without the cerebellum of clopidogrel-treated and placebo–treated APP23 mice were used. The hemispheres were homogenized 2 x 20 s at 6500 rpm (Precellys® 24, Bertin Technologies) in Tris-buffered saline (pH 7.6, 20 mM Tris, 137 mM NaCl, including protease inhibitors (Roche, Basel, Switzerland)). The homogenates were then sonicated and centrifuged at 100,000 × g for 1 h at 4°C. Supernatants (as TBS soluble fractions) were removed and stored at − 80 °C. The remaining pellets were resuspended in 1% Triton TX-100/TBS by sonication and then centrifuged at 100,000× g at for 1 h at 4°C. The supernatants were taken as membrane-bound Aβ and the pellets were resuspended in 2% SDS/TBS by sonication. After centrifugation at 100,000 x g at RT, the supernatants (protein-bound Aβ) were removed and the pellets were finally dissolved in 70% formic acid (FA). After the centrifugation at 100,000 × g for 1 h at 4°C,the FA-soluble supernatants (insoluble Aβ) were separated and neutralized with 1 M Tris (pH = 11.3, 1:20) and stored at − 80 °C. Total protein concentration was determined using Bio-Rad protein assay (Bio-Rad Protein Assay Kit II cat # 5000002). TBS- and FA-soluble levels of Aβ40 and Aβ42 were measured using sandwich-ELISAs from BioLegend (Legend MAX Human Amyloid Beta (1-42) and Amyloid Beta (1-40) ELISA kits) according to the manufacturer’s protocols and normalized to the total protein concentration.

### 2.6 Surface-Based Fluorescence Intensity Distribution Analysis Assay

Surface-based fluorescence intensity distribution analysis (sFIDA) assays were performed as described by Pils et al.(Pils et al., 2023). Briefly, capture antibody Nab228 (2.5 µg/ml in 0.1 M NaHCO₃) was added at 40 µl per well and incubated overnight at 4°C. Wells were washed five times with TBST and TBS. Blocking was done with 80 µl of 0.5% BSA in TBS-ProClin per well for 1.5 h at room temperature, followed by washing as described above. Samples, including soluble fraction diluted 1:10 and formic acid fraction diluted 1:50, were added at 20 µl per well in PBS-T with 0.5% BSA and 0.03% ProClin and incubated for 2 h at room temperature. Wells were washed again with five times TBS, then 20 µl per well of detection antibodies IC16 CF633 and Nab228 CF488A (both at 0.625 µg/ml in TBS-T with 0.1% BSA) were added and incubated for 1 h at room temperature. After a final wash with five times TBS, 80 µl of TBS-ProClin was added per well. Using TIRFM (Leica DMI6000B, Leica microsystems, Wetzlar, Germany), 3.15% of the well surface were imaged at 25 different positions (14-bit grayscale, channel 0: excitation: 635 nm, emission filter: 705 nm, exposure time: 1000 ms, gain: 1000, channel 1: excitation: 488 nm, emission filter: 525 nm, exposure time: 1000 ms, gain: 1000). Image data analysis was performed using the in-house software tool sFIDAta which enables automated detection and elimination of artefact-containing images and counting of co-localized pixels above a predefined cutoff value to reduce background noise (channel 0: 1500, channel 1: 1500). The calibration of pixel-based readouts into molar particle concentrations were performed as described by Pils et al. (Pils et al., 2023).

### 2.7 Western Blot

Brain homogenates in Tris-buffered saline were centrifuged at 10,000 × g for 10 min at 4°C. The supernatants were assayed for total protein (Precision Red Protein Assay Reagent, ADV02, Cytoskeleton). Equal amounts of protein were electrophoresed on an 8% or 12% SDS-PAGE gel under reducing conditions and transferred to a nitrocellulose blotting membrane (GE Healthcare Life Sciences). The membrane was blocked with 5% nonfat dry milk in PBS-T (PBS with 0.1% Tween 20) and then incubated with primary antibodies Iba1 (JM36-62, Invitrogen, 1:1000), GFAP (GA5, Invitrogen, 1:1000), GAPDH (14C10, Cell Signaling Technology, 1:2000) diluted in 5% BSA in PBS-T overnight at 4°C. After three washes with PBS-T, the membranes were incubated with HRP-conjugated secondary antibodies at 1:2500 dilution in 5% non-fat dry milk in PBS-T. Western Chemiluminescent HRP substrate solution (BioRad) and the Vilber Fusion-FX6-EDGE V.070 system and for quantification of the chemiluminescent signals Evolution-Capt EDGE software (Version 18, 02) were used for visualizing protein bands. The bands were quantified using ImageJ.

### 2.8 Flow cytometry

Heparinized blood from clopidogrel treated and placebo treated APP23 mice was diluted in Tyrode’s buffer (134 mM NaCl,12 mM NaHCO_3_, 2.9 mM KCl, 0.34 mM Na_2_HPO_4_, 20 mM HEPES, 10 mM MgCl_2_, 5 mM glucose, 0.2 mM CaCl_2_, pH 7.35) and washed twice. Blood samples were mixed with fluorophore-labeled antibodies from Emfret Analytics (P-selectin (Wug.E9-FITC) and JON/A-PE for an active form of αIIbβ3 integrin) after the addition of 2 mM CaCl_2_ and stimulated with the indicated agonists for 15 min at RT. For analysis of glycoprotein surface expression on platelets, antibodies from Emfret Analytics against GPVI (JAQ1-FITC), integrin β3 chain (GPIIIa, CD61, Luc.H11-FITC), GPIb (CD42, Xia.G5-PE), integrin ⍺5 (CD49e, Tap.A12-FITC) were added to the blood samples. The reaction was stopped by the addition of PBS and samples were analyzed on a FACSCalibur flow cytometer (BD Biosciences).

### 2.9 Statistical analysis

All statistical analyses were performed with GraphPad Prism (Prism 9; Graph Pad Software, Inc.). Unpaired t-test or one-way ANOVA, for two groups or three groups. The results were presented as the mean with each individual data point or in bar graph ± SEM. A p value <0.05 was considered significant (*p < 0.05, **p < 0.01).

## 3 Results

### 3.1 Increased permeability of the blood-brain barrier at late stages of Alzheimer pathology in mice

The dysfunction of the blood-brain barrier is a pathological hallmark of aging and Alzheimer’s disease (AD). This dysfunction facilitates the entry of blood-derived neurotoxins, inflammatory cells, and microbial pathogens into the brain, subsequently triggering neuroinflammation and neurodegeneration (Sweeney et al., 2018). To investigate whether the increase in platelet counts within the brain parenchyma was due to impaired BBB function, we assessed the integrity of the BBB by measuring blood-to-brain extravasation of immunoglobulin G (IgG). Figure 2 shows the abundance of IgG in the cerebral cortex (Fig. 1A) and the hippocampus (Fig. 1C) of WT; mT/mG;PF4Cre+ and APP23;mT/mG;PF4Cre+ mice at 6, 16 and 24 months of age. Quantification analysis revealed a significant but slight increase in IgG abundance in the cortex of 24-month-old control mice. However, the increase in IgG abundance was strongly elevated in APP23;mT/mG;PF4Cre+ mice at this age compared to 6- and 16-month-old mice, indicating BBB breakdown in aged mice. BBB breakdown was more pronounced in Alzheimer mice (Fig. 1B) and thus, a significant difference between WT;mT/mG;PF4Cre+ and APP23; mT/mG;PF4Cre+ mice was observed at the age of 24 month.

**Figure 1.**
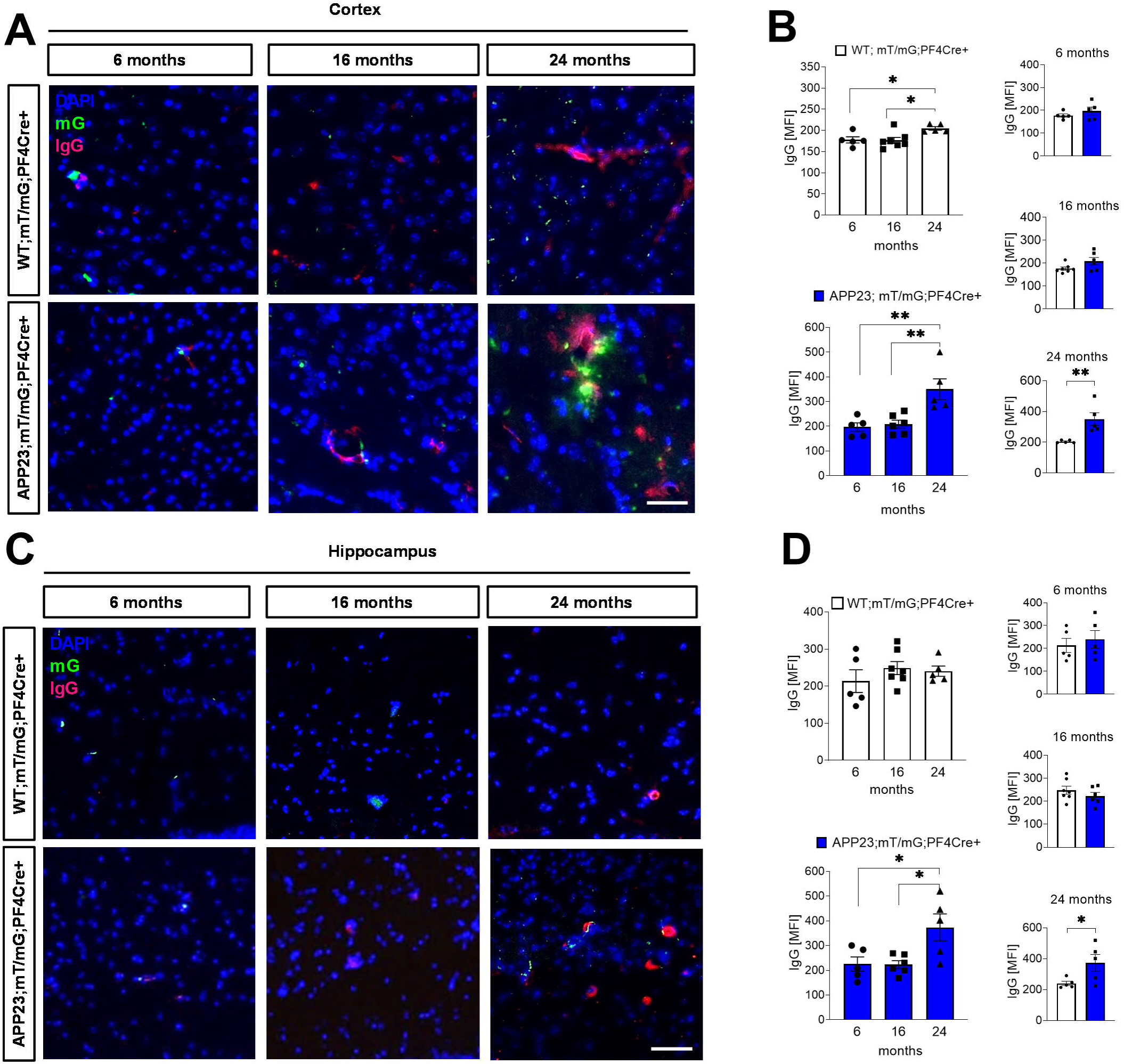
Increased permeability of the blood-brain barrier in the cerebral cortex and the hippocampus of APP23 mice. (A, C) Representative images of GFP expressing platelets (mG, green) and stained immunoglubolins G (IgG; pink) in the cerebral cortex and hippocampus regions of WT mT/mG;PF4Cre+ and APP23 mT/mG;PF4Cre+ mice at 6, 16 and 24 months of age. Cell nuclei were stained by DAPI (blue). Scale bar, 20 µm. (B, D) Quantification of IgG abundance in the cortex and hippocampus of WT;mT/mG;PF4Cre+ and APP23; mT/mG;PF4Cre+ mice at different age and comparison between the genotypes (n=5-7). Bar graphs indicate mean values ± SEM. Statistical analyses were performed using an ordinary one-way ANOVA followed by a Dunett’s post-hoc test or an unpaired t-test. *p < 0.05; **p < 0.01.

**Figure 2.**
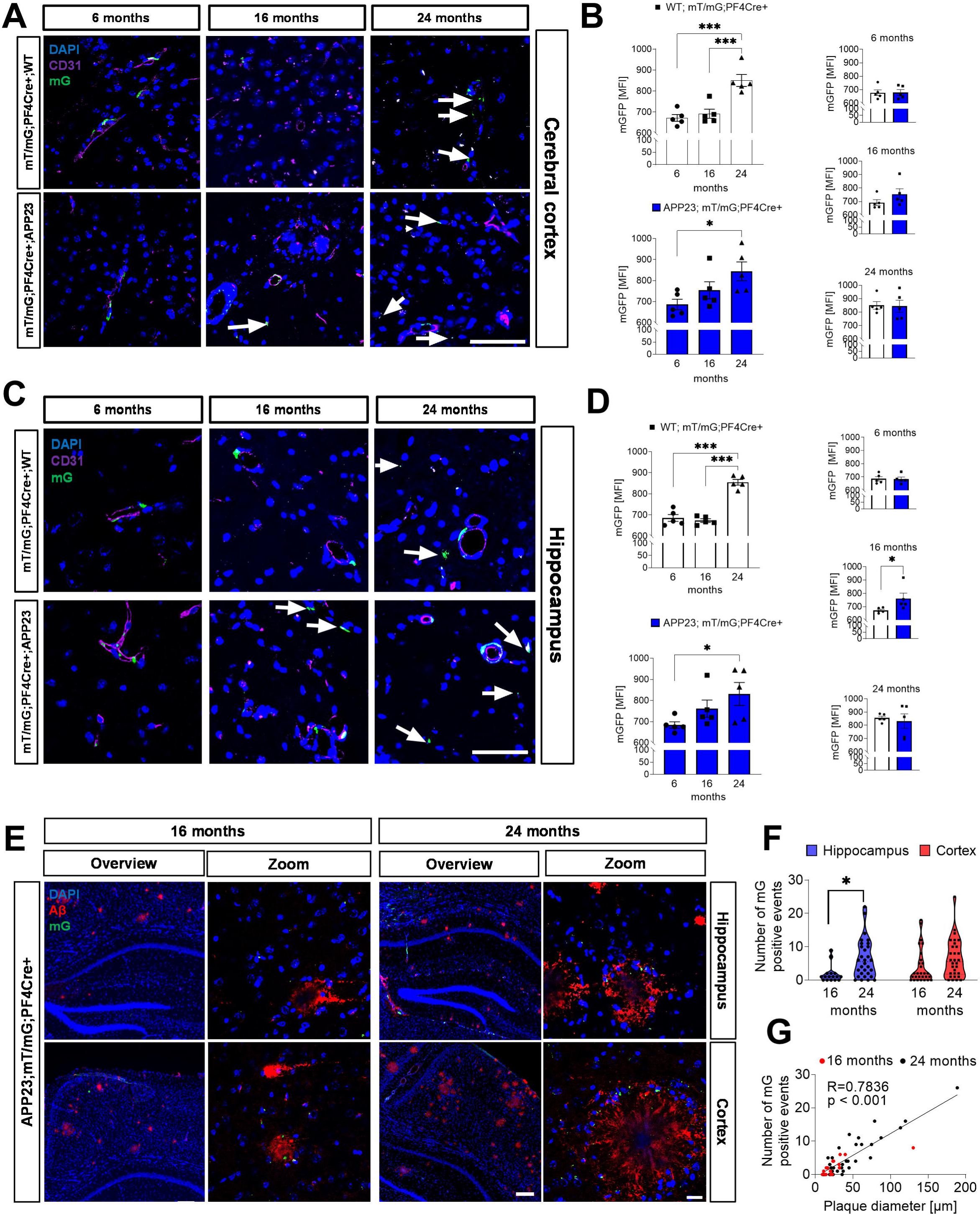
Early migration of platelets into the brain parenchyma and accumulation around amyloid plaques in transgenic APP23 mice. (A, C) Representative images of GFP-expressing platelets (mG, green) and stained endothelial cells (antibody CD31; purple) in the cerebral cortex and hippocampus regions of WT; mT/mG;PF4Cre+ and APP23; mT/mG;PF4Cre+ mice at 6, 16 and 24 months of age. Cell nuclei were stained by DAPI (blue). White arrows indicate platelets outside the vessels. Scale bar, 50 µm. (B, D) Quantification of platelet abundance in the cerebral cortex and the hippocampus of APP23 mT/mG;PF4Cre+ and WTmT/mG;PF4Cre+ mice. Mean fluorescence intensity (MFI) of mGFP was used for quantification. Statistical analyses were performed using an ordinary one-way ANOVA followed by a Dunett’s post-hoc test or an unpaired t-test. Bar graphs indicate mean values ± SEM, *p < 0.05; ***p < 0.001 (n=5). (E) Representative images of immunofluorescence staining of amyloid plaques using the 6E10 antibody (red) in the hippocampus (upper panel) and cerebral cortex (lower panel) of 16- and 24-month-old APP23mT/mG;PF4Cre+ mice. Overview scale bar, 200 µm. Zoom scale bar, 20 µm. (F, G) Quantification of the mGFP positive events around amyloid plaques in the hippocampus and cerebral cortex of 16- and 24-month-old APP23mT/mG;PF4Cre+ mice and spearmen correlation between the diameter of the amyloid plaques and the number of mG positive events (n=5 mice each with 3-9 plaques).

The quantification of the abundance of IgG in the hippocampal region did not show any significant differences in WT;mT/mG;PF4Cre+ mice at any age that has been analyzed (Fig. 1C and D). However, we observed a significant increase in IgG abundance in 24-month-old APP23;mT/mG;PF4Cre+ mice compared to 6- and 16-month-old APP23;mT/mG;PF4Cre+ mice. Thus, a significant difference of IgG abundance in brain parenchyma was observed in 24-month-old mice (Fig. 1D). In conclusion, APP23 mice showed an elevated increase of the permeability of the BBB in the cortex. In contrast, only aged transgenic APP23 mice showed an increase in BBB permeability in the hippocampus while control mice showed almost no increase in BBB permeability upon aging.

### 3.2 Platelets from APP23 mice enter the brain parenchyma at an earlier age than those from WT mice

The entry of blood cells, such as neutrophils, into the central nervous system has been described and observed in the brains of patients and mouse models of Alzheimer’s disease (Zenaro et al., 2015, Mapunda et al., 2022). However, it remains controversial to date whether or not blood platelets penetrate the brain (Kniewallner et al., 2015, Kniewallner et al., 2020). To analyze the transmigration of platelets through the blood brain barrier into the brain parenchyma, we crossed the platelet/megakaryocyte-specific mT/mG: PF4Cre^+^ reporter mouse, whose platelets express a GFP signal, with the transgenic APP23 mouse. The presence of platelets outside the blood vessels was analyzed in the region of the cerebral cortex (Fig. 2A and B) and the hippocampus (Fig. 2C and D) of WT mT/mG;PF4Cre+ and APP23mT/mG;PF4Cre+ mice at 6, 16 and 24 months of age. We observed a significant increase in platelets in the cortex of both, 24-month-old WT and APP23 mice compared to 6- and 16-month-old mice. There were no significant differences between WT and APP23 mice at any age (Fig. 2B). However, APP23 mice showed elevated platelet migration in to the brain parenchyma already at 16 month of age without reaching statistical significance. Quantitative analysis of platelets in the hippocampus showed that the abundance of platelets was significantly increased in APP23mT/mG;PF4Cre+ mice compared to WT mT/mG;PF4Cre+ mice at 16 months of age (Fig. 2D). However, platelet abundance was similar in both, WT and APP23 mice at 24 months of age, suggesting earlier platelet infiltration into the brain parenchyma in transgenic APP23 mice (Fig. 2D).

Previously, we provided evidence for platelet adhesion to vascular Aβ deposits that significantly contributes to the aggregation of Aβ leading to CAA (Gowert et al., 2014, Donner et al., 2016). Therefore, we investigated whether platelets also accumulate in areas of Aβ plaques in the brain parenchyma using 16 and 24-month-old APP23 mice with pronounced Aβ pathology. The staining of Aβ and quantitative analyses of the hippocampus and the cerebral cortex revealed an accumulation of platelets around amyloid plaques (Figure 2E). Furthermore, there was a positive correlation between the diameter of amyloid plaques and an increased number of platelets or platelet-aggregates in the brain parenchyma (Figure 2E-G).

### 3.3 Platelets co-localize to microglia in different brain regions of APP23 mice

Microglia function as resident phagocytes that dynamically monitor the environment and play a critical role in CNS tissue maintenance, response to injury, and defense against pathogens. Amyloid plaques surrounded by activated microglia are part of the inflammatory processes in AD brains. In addition, microglia are already known to be important for Aβ uptake and degradation. However, continuous activation of microglia by Aβ aggregates or fibrils leads to microglial production of TNFα, IL-1β and other pro-inflammatory cytokines, that may contribute to neurodegeneration (Hansen et al., 2018). In the present study, immunofluorescence was used to investigate the localization and activation of microglia by IBA-1 staining in relation to platelet localization. Activation of microglia is associated with increased IBA-1 expression, which may be involved in membrane ruffling and phagocytosis. However, it is considered as a marker for all microglia, as non-activated microglia also show IBA-1 protein expression (Hopperton et al., 2018). As shown in Figure 3, *APP23mT/mG;PF4Cre+* mice display high numbers of IBA-1-positive microglia. High abundance of IBA-1 positive microglia were present in the hippocampus and the cerebral cortex of 24-month-old *APP23mT/mG;PF4Cre+* mice. To a lesser extent, IBA-1 positive microglia were found in 16-month-old *APP23mT/mG;PF4Cre+* mice. In contrast, only low levels of IBA-1-positive microglia were detected in *WTmT/mG;PF4Cre+* mice at 16 and 24 months of age. IBA-1-positive cells are distributed throughout the parenchyma in *WTmT/mG;PF4Cre+* mice. In contrast, IBA-1-positive cells are concentrated in specific regions, most likely around amyloid plaques in *APP23mT/mG;PF4Cre+* mice. Interestingly, in 24 months old *APP23mT/mG;PF4Cre+* mice, platelet positive fluorescent signals directly merge with IBA-1 microglia staining as shown in Figure 3. This indicates a close proximity of the two cell types in AD transgenic mice.

**Figure 3.**
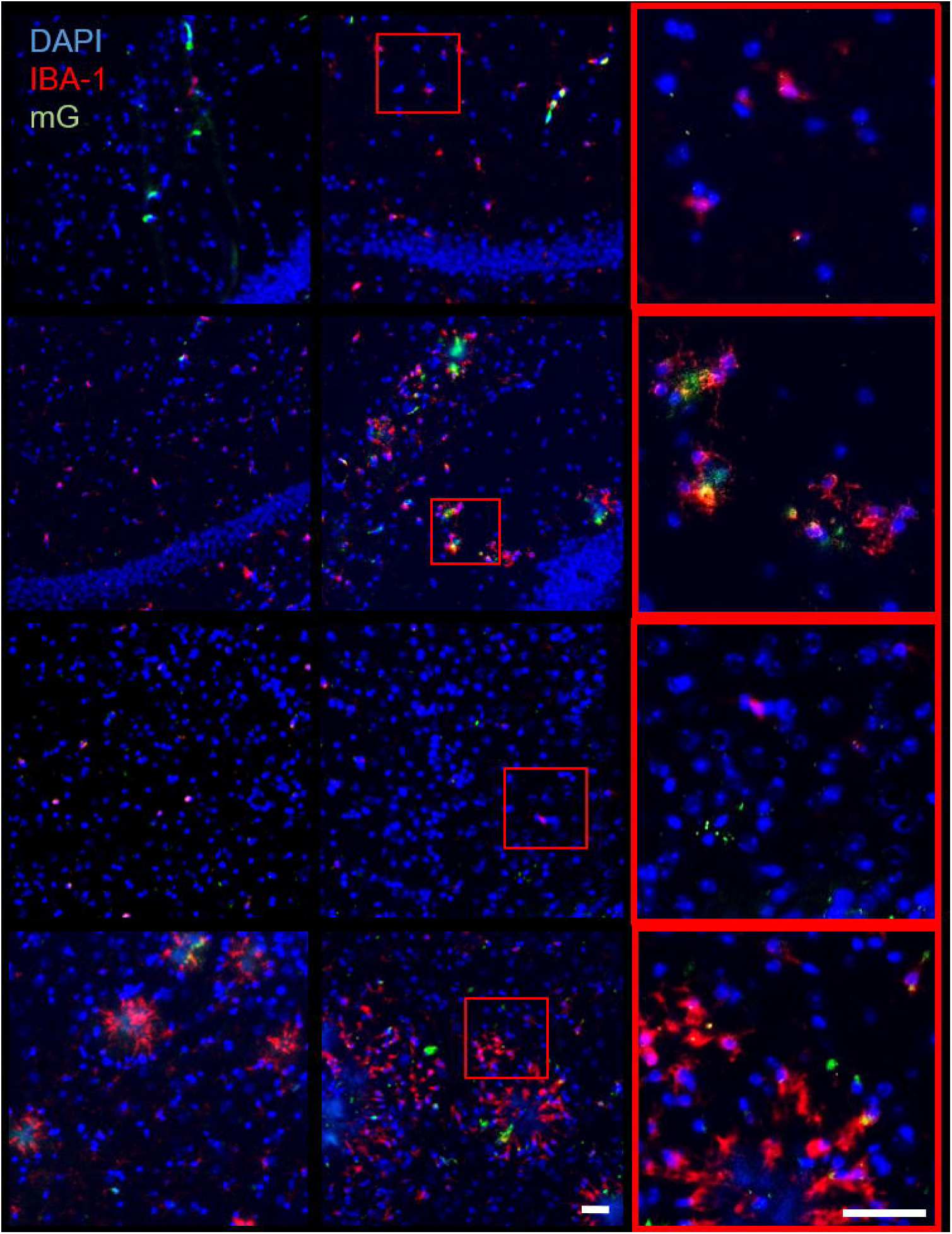
Analysis of platelet-microglia interaction in different brain regions of APP23mT/mG;PF4Cre+ and WTmT/mG;PF4Cre+ mice at different time points. (A) Representative images of immune fluorescent staining of microglia (IBA-1, red) in the hippocampus and the cerebral cortex of APP23mT/mG;PF4Cre+ and WT control mice. Platelets express a GFP signal (mGFP, green). Cell nuclei were stained with DAPI (blue). Scale bar 50 µm, n=4-5.

### 3.4 Platelets are in close proximity to neurons in the hippocampus and cortex of APP23 mice

Neuronal loss is a late feature of Alzheimer’s disease. In *APP23mT/mG;PF4Cre+* mice, neuronal loss start with 14 -18 months of age and worsens over time (Calhoun et al., 1998). To determine whether or not platelets are located near neurons, neuronal nuclei in 16 and 24-month-old *WTmT/mG;PF4Cre+* and *APP23mT/mG;PF4Cre+* mice were stained with an antibody against neuronal nuclear protein (NeuN; purple) and the hippocampus and the cerebral cortex were analyzed. As shown in Figure 4, in 16 month old *WTmT/mG;PF4Cre+* mice, almost no platelets were observed near to neuronal nuclei. In contrast, in 16 months old *APP23mT/mG;PF4Cre+* mice and in 24-month-old *WTmT/mG;PF4Cre+* and *APP23mT/mG;PF4Cre+* mice, platelets are partially located near to neurons in the hippocampus (Figure 4A) as well as in the cerebral cortex (Figure 4B).

**Figure 4.**
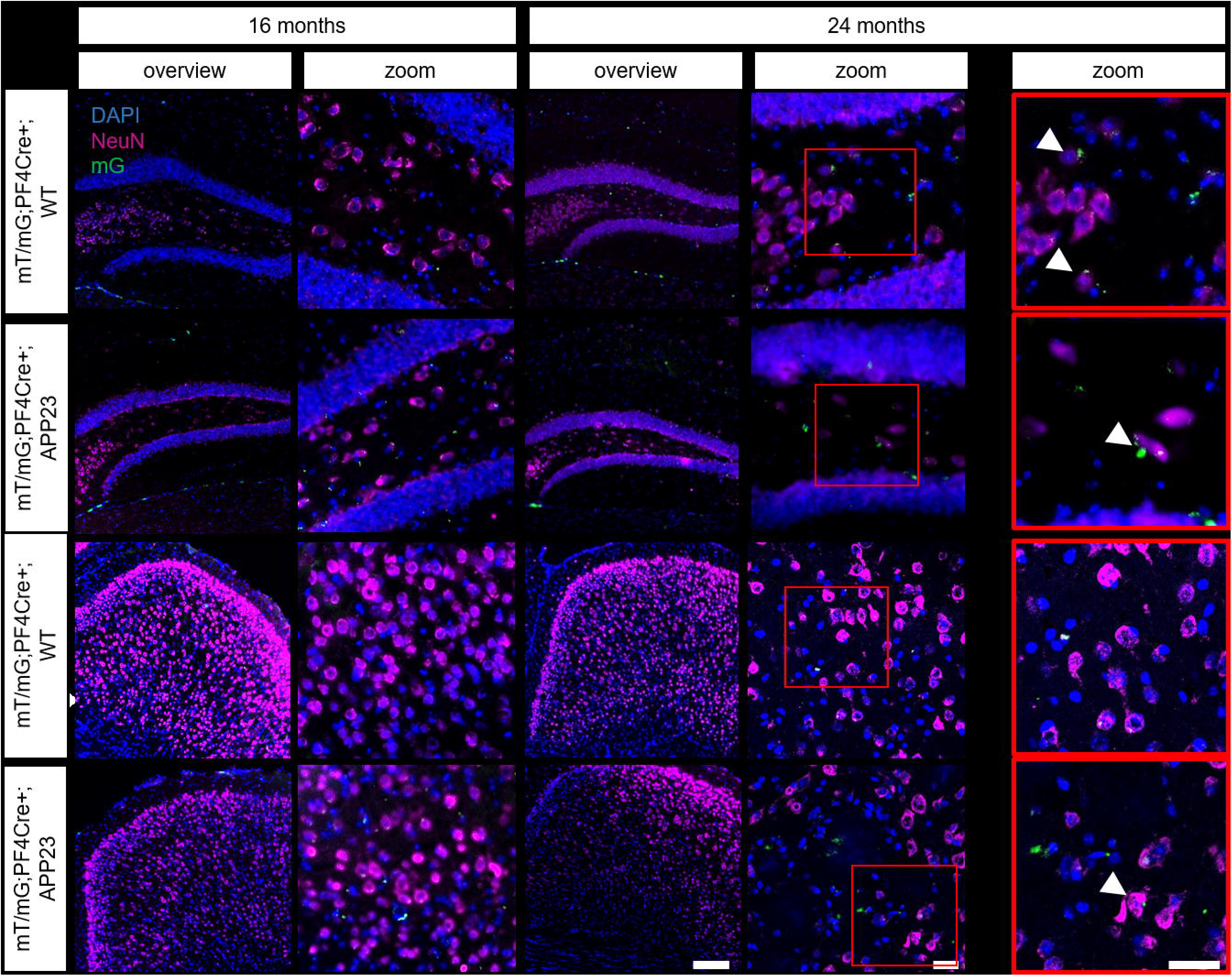
Analysis of platelet-neuron interaction in different brain regions of APP23mT/mG;PF4Cre+ and WTmT/mG;PF4Cre+ mice at the age of 6 and 24 month of age. Representative images of immune fluorescent staining of neuronal nuclear protein (NeuN; purple) to detect neurons in the (A) hippocampus and (B) in the cerebral cortex of 16 and 24 month old APP23mT/mG;PF4Cre+ and WT control mice. Platelets express a GFP signal (mGFP, green). Cell nuclei were stained with DAPI (blue). Scar bars: Overview 200 µ, zoom-in 20 µm, n=3.

### 3.5 Neurodegeneration and gliosis in APP23 mice is not altered by long-term antiplatelet treatment with clopidogrel

Neuronal loss is a pathological hallmark of Alzheimer’s disease (AD). The APP23 mouse model has been shown to exhibit neuronal loss in the CA1 region of the hippocampus in 14-18-month-old mice (Calhoun et al., 1998). Moreover, we here provide evidence that platelets localize near to microglia (Figure 3) and neurons (Figure 4) in the hippocampus as well as in the cerebral cortex of APP23 mice.

To investigate the effects of anti-platelet therapy by clopidogrel treatment on neurodegeneration in the hippocampus, we performed NeuN immunostaining and counted the NeuN/DAPI-positive cells in the CA1 region (Fig. 5A and B). The results showed no significant differences between the clopidogrel and placebo groups, indicating that antiplatelet treatment did not affect neurodegeneration in the hippocampus (Fig. 5B). In addition, NeuN levels in brain lysates were also determined by western blot analysis but showed no significant differences between clopidogrel and placebo treated APP23 mice (Fig. 5C and D).

**Figure 5.**
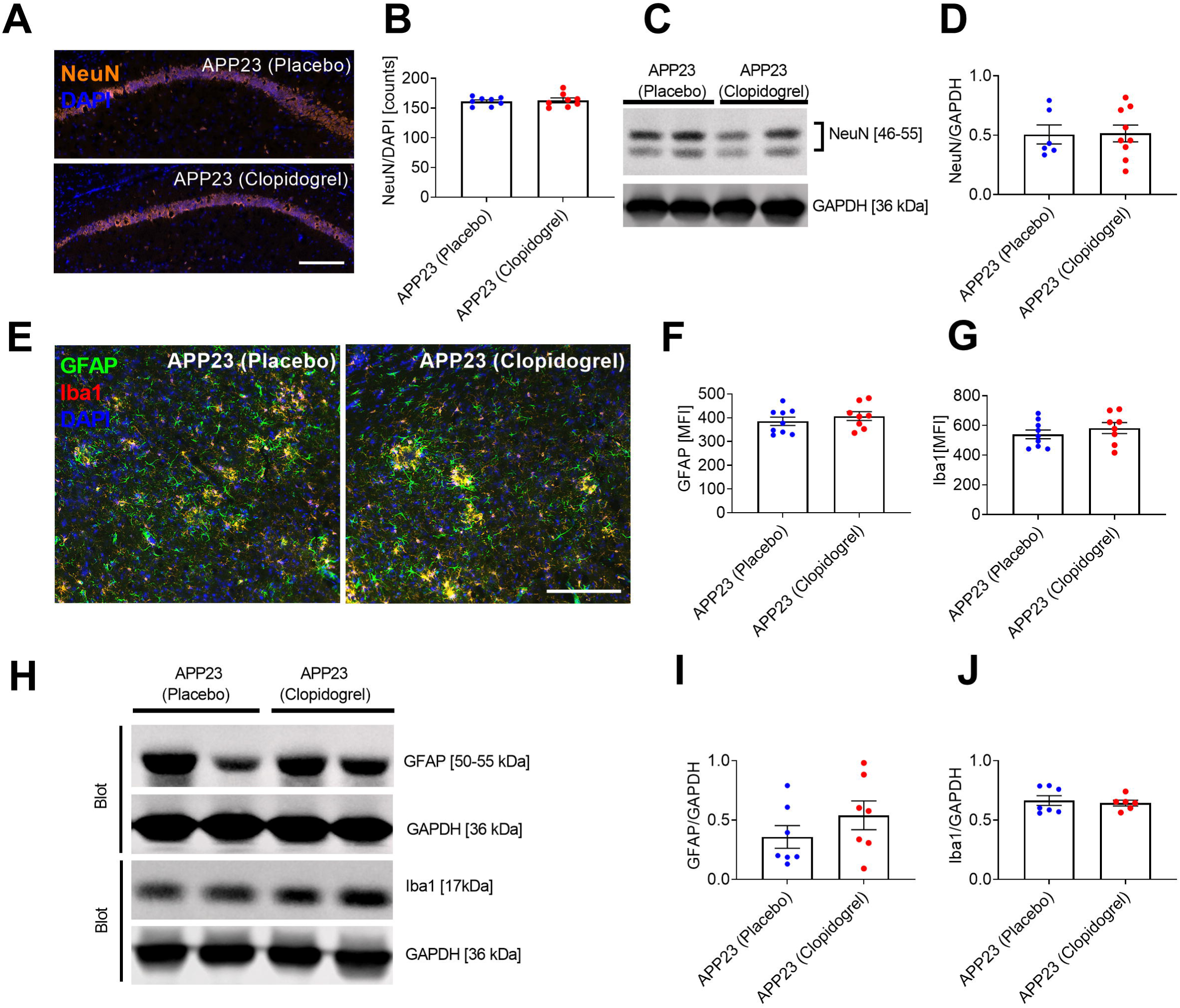
Unaltered neurodegeneration and gliosis in the brains of APP23 mice after treatment with clopidogrel. (A, B) Representative images of NeuN and DAPI staining and quantification of the number of NeuN/DAPI positive cells in the CA1 hippocampal region. Scale bar, 200 µm. (C) NeuN levels in whole brain lysates were detected by western blot analysis. GAPDH was used as loading control. (D) NeuN levels were quantified by densitometric measurement. (E) Immunofluorescence detection of Iba1-positive activated microglia (red) and GFAP-positive activated astrocytes (green). Scale bar, 200 µm. (F, G) Quantification of GFAP and Iba1 are mean intensities. (H) Levels of GFAP and Iba1 in whole brain lysates were detected by western blot analysis. GAPDH was used as a loading control. (I, J) Levels of GFAP and Iba1 were quantified by densitometric measurement. Bar graphs indicate mean values ± SEM. Statistical analyses were performed using an unpaired t-test.

To evaluate the effect of clopidogrel treatment on disease-associated gliosis as a marker of neuroinflammation, we analysed the brains of APP23 mice for reactive astrocytes using glial fibrillary acidic protein (GFAP) and for microglia using Iba1 (Fig. 5E). No differences in GFAP (Fig. 5E and F) and Iba1 (Fig. 5E and G) signal intensity were observed between placebo and clopidogrel treated APP23 mice. To confirm these findings, we performed western blot analysis on total brain protein extracts from mice (Fig. 5H). Analysis of GFAP and Iba1 protein levels showed no statistically significant differences between the groups, indicating that gliosis was not affected by clopidogrel treatment of APP23 mice (Fig. 5I and J). In conclusion, long-term treatment with clopidogrel has no effect on neurodegeneration and gliosis in APP23 mice.

### 3.6 Long-term antiplatelet treatment with clopidogrel reduces CAA and soluble Aβ42 levels but does not impact parenchymal amyloid plaque burden

In our previous study, we treated 13-month-old transgenic APP23 mice, which already had parenchymal plaques and were beginning to develop vascular plaques, for three months, and observed a significant reduction in vascular amyloid plaques compared to untreated APP23 mice (Donner et al., 2016). To investigate the effects of long-term clopidogrel treatment, we started the treatment in APP23 mice at 2 months of age and treated the mice for 15 months. At the end of the treatment period, we analyzed the effect of long-term clopidogrel treatment on blood parameters. The treatment did not cause any changes in blood cell counts or platelet volume (Fig. S1A-C) or in the expression of glycoproteins on the surface of platelets (Fig. S1E and F). Measurements of activated ⍺IIbβ3 integrin and P-selectin exposure after platelet stimulation using flow cytometry confirmed the inhibitory effect of clopidogrel on platelets. (Fig. S1G and H).

Immunohistochemical and biochemical experiments were subsequently performed to investigate the impact of long-term clopidogrel treatment on Aβ pathology, neuroinflammation, and neurodegeneration. The staining of brain tissue sections with an Aβ antibody (6E10) was performed to investigate the impact of long-term clopidogrel treatment on Aβ pathology in APP23 transgenic mice. Figure 6 shows the quantification of amyloid plaques in the cerebral cortex and in the hippocampus (Fig. 6A - C). Immunohistochemical analysis revealed no significant differences in Aβ plaque load or plaque numbers in the brain parenchyma of the cortex (Fig. 6B) and the hippocampus (Fig. 6C) between placebo and clopidogrel treated APP23 mice. Additionally, we quantified Aβ40 and Aβ42 biochemically in the TBS fraction (soluble Aβ) and in the formic acid fraction (insoluble Aβ) of brain homogenate extracts (Fig. 6D). No significant differences were found in the Aβ40 content of any of the fractions between the clopidogrel and the placebo treated mice (Fig. 6D). However, we detected a significant reduction in the concentration of soluble Aβ42 in the TBS fraction (Fig. 6D). The concentration of Aβ oligomers in both, the TBS and the formic acid fraction was analysed using the surface-based fluorescence intensity distribution analysis (sFIDA) assay (Kulawik et al., 2018). No significant differences in oligomeric Aβ concentrations were found between the groups in the TBS fraction (Fig. 6E). However, an increased concentration of oligomeric Aβ was detected in the plaque-associated fraction (formic acid fraction) of brain lysates from clopidogrel treated compared to untreated APP23 mice (Figure 6E). Consistent with our previous study (Donner et al., 2016), we observed a significant decrease in CAA following treatment with clopidogrel (Fig. 6F). Since CAA is associated with brain hemorrhages, we performed Prussian blue staining to detect the cerebral microhaemorrhages. No differences in the number of microhaemorrhages were observed between placebo and clopidogrel treated APP23 mice, suggesting that clopidogrel did not have a negative effect on hemorrhages at this age (Fig. S2 A and B). In conclusion, long-term treatment with clopidogrel leads to no reduction of Aβ burden in the brain parenchyma, but affects CAA in brain vessels and the soluble Aβ42 levels in the brain of APP23 mice.

**Figure 6.**
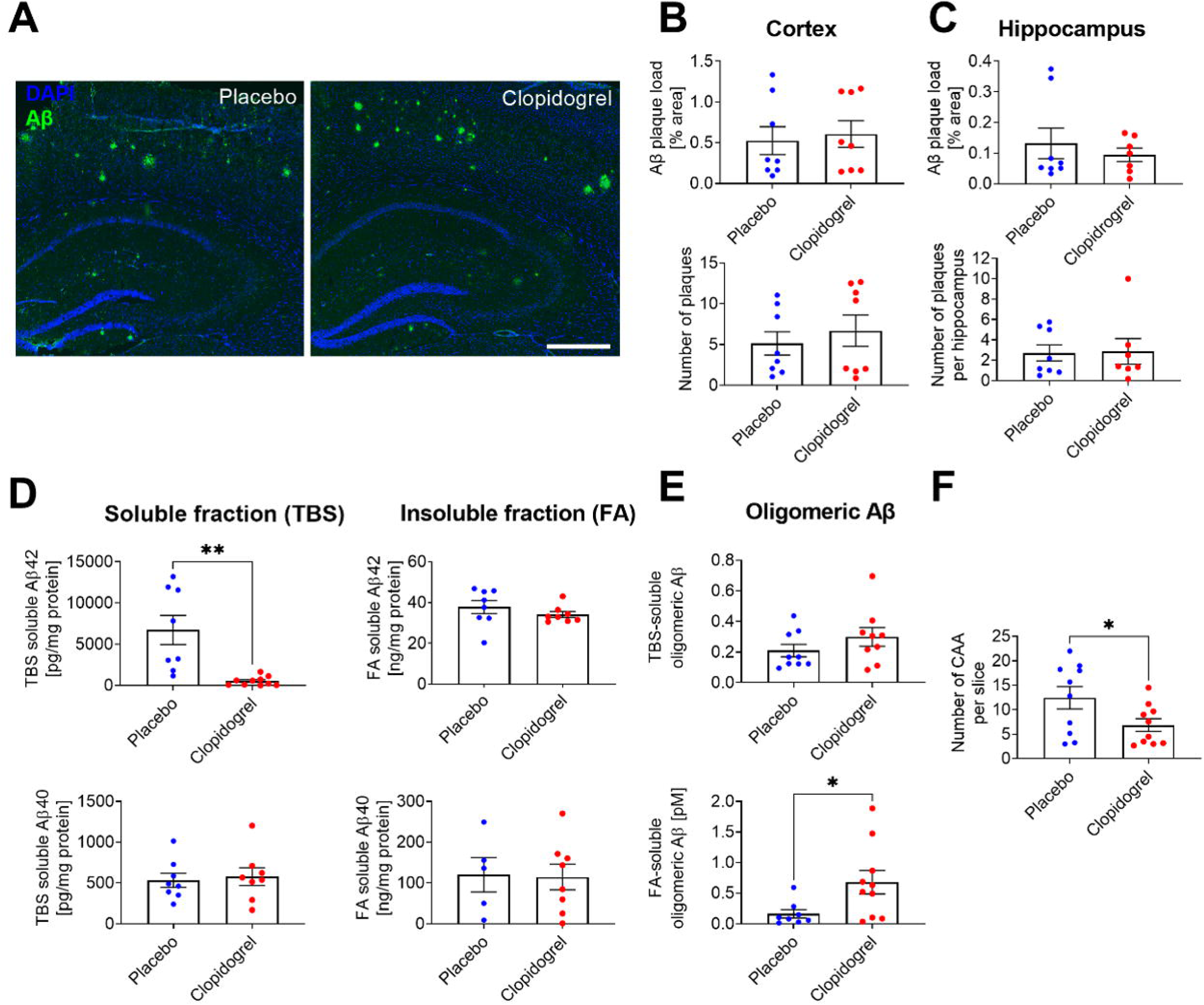
Effects of clopidogrel treatment on Aβ pathology in APP23 mice. (A) Representative images of Aβ plaques in the hippocampus and cortex of untreated (placebo) and clopidogrel treated APP23 mice. Amyloid plaques were visualised by immunostaining with the Aβ antibody 6E10. Scale bar: 500 µm. (B, C) Quantification of the percentage of cortical and hippocampal areas occupied by plaques and the number of plaques. (D) Levels of Aβ40 and Aβ42 in Tris-buffered saline (TBS) and formic acid (FA) fractions of brain homogenates from clopidogrel treated and untreated (placebo) APP23 mice were analysed by ELISA. (E) Aβ oligomer concentrations in the above fractions were analysed by sFIDA assay. (F) Cerebral amyloid angiopathy (CAA) was detected by congo red staining and the number of amyloid-affected vessels was quantified. Scale bar: 100 µm. (G, H) Aβ40-positive vessels were detected using an anti-Aβ40 antibody, and the number and area were quantified. Scale bar: 50 µm. (I, J) Aβ42-positive vessels were detected using an anti-Aβ42 antibody, and the number and area were quantified. Scale bar: 50 µm. Bar graphs indicate mean values ± SEM. Statistical analyses were performed using an unpaired t-test. *p < 0.05; **p < 0.01.

## 4 Discussion

AD is a multifactorial disorder in which various pathophysiological events contribute pathogenesis and progression (Selkoe, 2011, Li and Selkoe, 2020). Platelets have been suggested to play a role in the pathogenesis of both AD and CAA (Stellos et al., 2010, Slachevsky et al., 2017, Donner et al., 2016, Jarre et al., 2014). They are important source of Aβ peptides and can release Aβ peptides following platelet activation (Gowert et al., 2014, Chen et al., 1995, Li et al., 1998). Moreover, the dysfunction of platelets is associated with AD and could be an initial contributor to AD pathology (Sevush et al., 1998, Johnston et al., 2008, Stellos et al., 2010, Slachevsky et al., 2017).

There are controversial studies on whether platelets are able to migrate into brain parenchyma in AD. In one study, platelet migration was not detected both in the APP-PS1 mouse model of Alzheimer’s disease, which develops amyloid plaques in the brain parenchyma but no CAA pathology, and in brain slices from Alzheimer’s patients (Kniewallner et al., 2015). In another study, platelets were detected outside the vessels in the brain parenchyma using immunofluorescence staining in the same mouse model APP-PS1 (Kniewallner et al., 2020). Here, using a reporter mouse model in which platelets express the GFP signal, we observed the same abundance of platelets in the brain parenchyma in both in the aged APP23 AD mouse model and in age-matched WT mice, suggesting an age-related process of platelet migration (Fig. 1). However, platelet burden in the hippocampal region was significantly higher in 16-month-old APP23 mice compared to WT mice. This result indicates an earlier onset of platelet entry into the hippocampal region in the AD mouse model. Whether platelet migration into the brain parenchyma is an active or passive process remains to be clarified. Our analysis of BBB leakage showed that the significant difference in BBB permeability between WT and APP23 mice did not appear until 24 months of age and not earlier. Already in other pathological conditions, such as multiple sclerosis (Kocovski et al., 2019) and stroke (Schleicher et al., 2015), it has been shown that platelets infiltrate into the brain parenchyma and affect neurodegeneration.

Previously, we showed that short-term (3 months) treatment with clopidogrel leads to the reduction of Aβ plaques in cerebral vessels (CAA) of transgenic AD model mice (Donner et al., 2016). The present study demonstrated that initiating clopidogrel treatment at an earlier stage prior to the onset of cognitive deficits and Aβ pathology and maintaining treatment for a prolonged time period of 15 month resulted, not only reduced CAA, but also reduced soluble Aβ42 levels. Although, a reduction in CAA and soluble Aβ42 level in the brain as well as an increase in plaque-associated oligomeric Aβ was observed in clopidogrel treated APP23 compared to control mice, we did not detect alterations in Aβ burden and gliosis in the brain parenchyma.

Previously, the effect of clopidogrel was investigated in an AlCl_3_-induced rat model of AD in which changes in neuroinflammatory mediators and proinflammatory cytokines, cholinergic dysfunction and oxidative stress lead to neurodegeneration (Khalaf et al., 2020, Akhtar et al., 2022). This is the preferred model for prophylactic treatment of Alzheimer’s disease (Akhtar et al., 2022). The authors observed that clopidogrel could alleviate learning and memory deficits in rats and suggested that the neuroprotective effect of clopidogrel was due to its anti-inflammatory effect (Khalaf et al., 2020). In the present study, we used a transgenic mouse model with stable sevenfold overexpression of mutant human APP carrying the pathogenic Swedish mutation (Sturchler-Pierrat et al., 1997). The latest study shows that this mouse model develops Aβ fibrils similar to the human type II fold, and is therefore a suitable model for the familial form of Alzheimer’s disease (FAD) (Zielinski et al., 2023). Although FAD and sporadic

AD may share some similarities, there are some important differences in manifestation of pathology and underlying cause, which could explain why we did not detect any changes in plaque burden in the parenchyma, neuroinflammation (gliosis) or neurodegeneration in clopidogrel treated APP23 mice compared to untreated APP23 mice. Previous behavioral study showed that APP23 mice exhibit severe deficits in spatial memory before amyloid plaques form, and that these initial deficits are related to the level of soluble Aβ42 in the brain (Van Dam et al., 2003). In this study, we detected strong reduced soluble Aβ42, but not of plaque-associated Aβ42, level in the brain from clopidogrel treated APP23 mice. This is consistent with the well-known fact that plaque burden does not correlate with cognitive decline in humans (Arriagada et al., 1992, Samuel et al., 1994). Although plaque burden in the brain parenchyma was unchanged by clopidogrel treatment, we detected an increase in oligomeric Aβ in the plaque-associated fraction but not in the soluble fraction. In recent years, there has been increasing evidence that soluble Aβ oligomers, much more than fibrilar amyloid plaque cores or Aβ monomers, are neurotoxic, disrupt synaptic function and contribute to cognitive decline in AD (Li et al., 2018, Hampel et al., 2021). There is also evidence that amyloid plaques can sequester the toxic Aβ oligomers and prevent their neurotoxic effects (Shankar et al., 2008). On the other hand, amyloid plaques are thought to act as a local reservoir for soluble oligomers (Koffie et al., 2009). Moreover, it is crucial to highlight that the measured concentration of oligomers, both in the soluble (TBS) and plaque-associated (FA) fractions, is exceedingly low and may not be a significant contributor to pathology in this mouse model.

Consistent with our previous study (Donner et al., 2016), the antiplatelet treatment with clopidogrel led to a significant reduction of CAA in APP23 mice. CAA is associated with disruption of vascular structure and hemorrhages contributing to cognitive impairment in AD (Smith, 2018). On the other hand, CAA causes cognitive decline independently of AD and is associated with depression, anxiety and apathy (Smith et al., 2021). Previously, we have shown that platelet recruitment to vascular Aβ deposits can lead to full occlusion of the vessel in APP23 transgenic mice (Gowert et al., 2014). Thus, hypo-perfusion in CAA affected cerebral vessels might be strongly reduced in clopidogrel treated APP23 mice leading to improved delivery of oxygen and nutrients to brain tissue. In addition, the P2Y_12_ mediated inhibition of ADP signaling of platelets may restrict or even avoid the release of endogenous murine Aβ peptides from activated platelets since we have already shown that ADP plays an important role in platelet mediated amyloid aggregation and CAA (Donner et al., 2016). Furthermore, using APP23 mice on an App-null background, Mahler et al. showed that endogenous murine Aβ increases cerebral Aβ load [52]. Another explanation for the reduction in CAA could be that the amount of fibrinogen is reduced in CAA. In particular, fibrinogen has been discussed for years as a cerebrovascular risk factor for AD, and identified fibrinogen as a contributing factor to CAA pathogenesis (Strickland, 2018, Cortes-Canteli et al., 2015, Paul et al., 2007). In our recent study, we showed that the binding of Aβ to the GPVI receptor on platelets leads to the release of fibrinogen, which can bind to Aβ peptides and is incorporated into amyloid aggregates (Donner et al., 2020, Donner et al., 2016).

This study has some limitations. One limitation is the use of only one sex (male). As there are sex-specific differences in pathologies in the APP23 Alzheimer’s mouse model, therefore, future analyses with female mice are needed. Another limitation is that we’ve only used a mouse model of AD that reflects familial AD and CAA, like all existing mouse models of AD but not sporadic AD, which is the most common form in humans and a complex multifactorial disease. In particular, it would be interesting to know whether antiplatelet therapy would affect the pathology in the sporadic mouse models of Alzheimer’s disease.

In conclusion, long-term antiplatelet treatment just before the onset of Aβ pathology in transgenic APP23 mice had no effect on neuroinflammation, neurodegeneration and Aβ plaques in the brain parenchyma in transgenic APP23 mice. However, we detected reduced soluble Aβ42 levels in the brain and reduced CAA. Further *in vivo* studies are needed to further investigate the potential benefits and risks of antiplatelet therapy in Alzheimer’s disease.

## 6 Author contribution

ME designed the study. LD, LMT, MP, FR, and OB performed experiments. LD, OB and ME analyzed and interpreted data. LD and ME wrote the manuscript with all authors providing feedback.

## 8 Acknowledgements

We thank Martina Spelleken for providing outstanding technical assistance.

## 10 Data Availability Statement

The original contributions presented in the study are included in the article, further inquiries can be directed to the corresponding author.

