## Supplementary Material for "Long-term antiplatelet therapy protects against cerebral but not parenchymal amyloid plaque formation and neurodegeneration in transgenic mice of Alzheimer’s disease"

### 1 Supplementary Figures and Tables

#### 1.1 Supplementary Figures

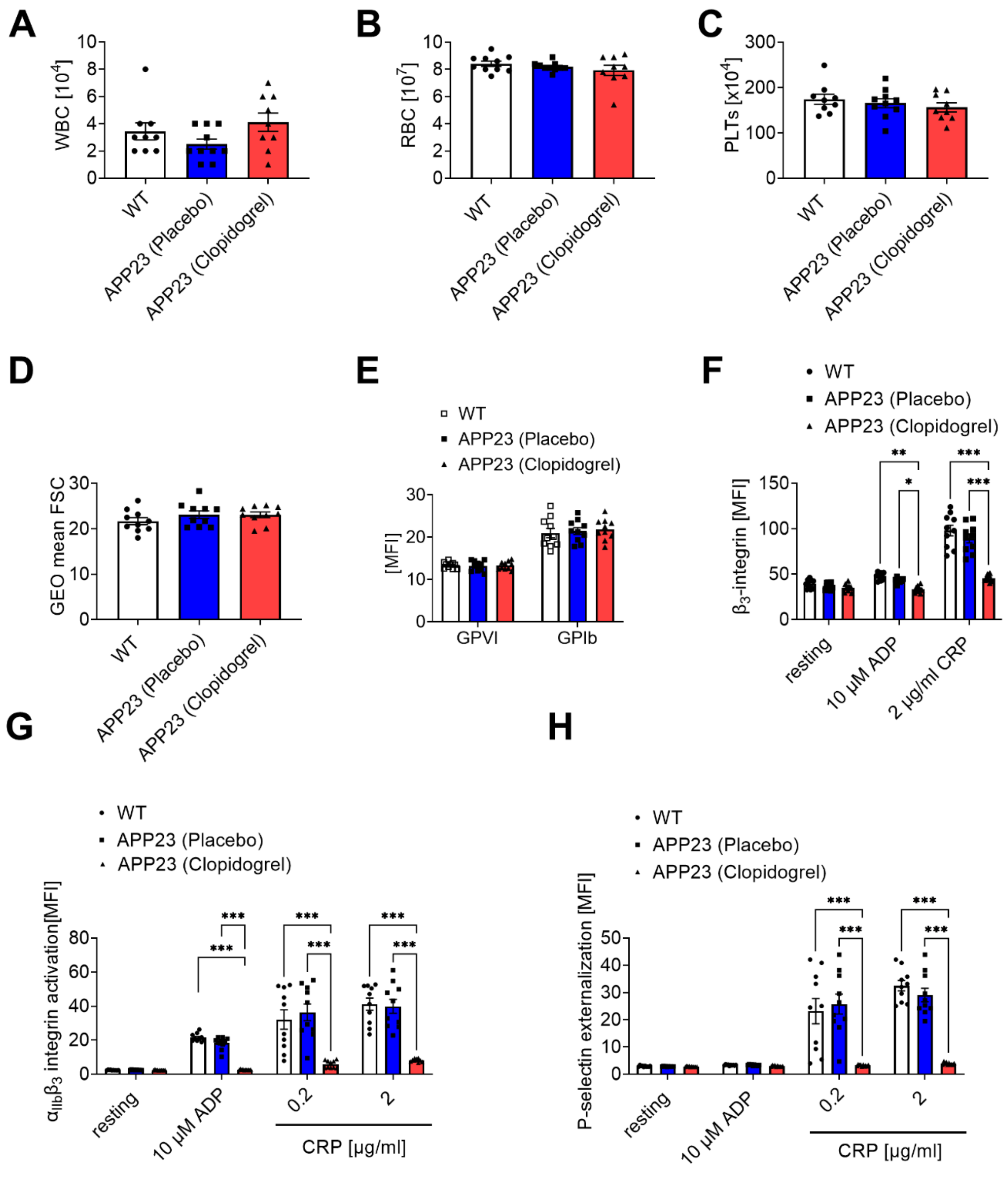


**Supplementary Figure 1.** Analysis of WT, placebo and clopidogrel treated APP23 mice at the end of the treatment. (A-C) Unaltered white blood cell (WBC), red blood cell (RBC) and platelet (PLTs) counts (n=9-10). (D) Platelet size measured by geometrical mean (n=10). (E) Expression of glycoprotein GPVI and GPIb on the surface of platelets and (F) externalization of the β3 integrin subunit upon platelet stimulation measured by MFI in flow cytometric analysis (n=10). (G, H) Significantly reduced αIIbβ3 integrin activation and P-selectin externalisation on the platelet surface of clopidogrel treated APP23 mice after stimulation (n=10). MFI: mean fluorescence intensity; CRP: collagen-related peptide. Bar graphs indicate mean values ± SEM. Statistical analyses were performed using one-way-Anova or two-way-Anova. *p < 0.05; **p < 0.01, ***p < 0.001.

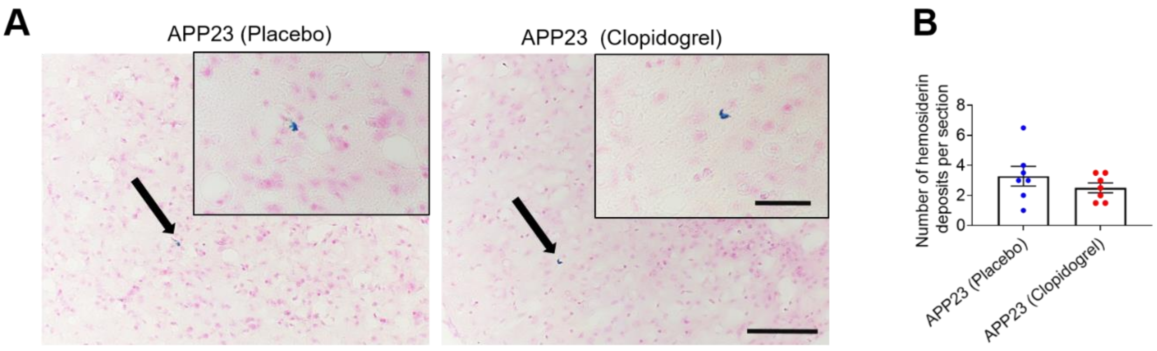


**Supplemental Figure 2.** No differences of cerebral micro hemorrhages between placebo and Clopidogrel treated APP23 mice. (A) Histological sections stained with Perls` Prussian blue indicate the presence of hemosiderin deposits. Scale bar in low magnification, 50 µm; in high magnification, 20 µm. (B) The number of haemosiderin deposits was quantified.
